# Entangling snakebite dynamics: the spatiotemporal role of rainfall on snake envenoming in Colombia

**DOI:** 10.1101/2021.07.13.452215

**Authors:** Carlos Bravo-Vega, Mauricio Santos-Vega, Juan Manuel Cordovez

## Abstract

The role of climate forcing on the population dynamics of infectious diseases has typically been addressed via retrospective analyses of aggregated incidence records over whole political regions. A central question in epidemiology has been whether seasonal and interannual cycles are driven by climate variation, or instead generated by other factors such as poverty or underreporting. Here, we use process-based models to determine the role of rainfall in the dynamics of snakebite, which is one of the main neglected tropical diseases around the world. We combined space-time datasets of snakebite incidence and rainfall for Colombia in combination with stochastic epidemiological models and iterated filtering methods to show the response to rainfall forcing, specifically, modulating the encounter frequency with venomous snakes. We identified six zones with different rainfall patterns to demonstrate that the relationship between rainfall and snakebite incidence was heterogeneous. Rainfall only drives snakebite incidence in regions with marked dry seasons, where rainfall becomes the limiting resource. In addition, the encounter frequency also differs between regions, and it is higher in regions where *Bothrops atrox* can be found. Our results show how the heterogeneous spatial distribution of snakebite risk seasonality in the country may be related to important traits of venomous snakes’ natural history.

**SIGNIFICANCE STATEMENT:** The association between seasonal climatic variables and diseases’ epidemiology has helped to understand disease burden under changing environments. For example, for several tropical zoonotic diseases rainfall has been identified as a critical covariate inducing incidence seasonality. Snakebite envenoming is a disease that affects mostly economically deprived populations, and the availability of treatment is scarce. However, the role of environmental factors on snakebite is still missing in the literature. We formulated an epidemiological model to quantify the role of rainfall on snakebite seasonality throughout Colombia. We found that rainfall has a significant effect on incidence in places with a marked dry season (Caribbean and Orinoco plains), but in areas without dry season (Amazonian basin and southwestern Colombia) incidence exhibits no seasonality. This study is the first epidemiological modeling approach to snakebite and underscores the importance of rainfall as the limiting resource in this system. Thus, it is important to consider the interaction between climate forcing and venomous snakes’ ecology as determinants of envenomation risk.

## INTRODUCTION

Snakebite envenoming is the neglected tropical disease (NTD) with the highest mortality rate around the world, but despite its importance data collection is still challenging and the real disease burden is underestimated (1, 2). Currently, the most reliable estimates are at least 1.8 million cases with 435.000 deaths each year, but this data can underestimate the real burden because patients prefer seeking traditional medicine instead of medical care (3–8). To perform estimations of real burden, it is important to understand the ecological characteristics that modulate the likelihood of human-snakes encounters, where the climate can play an important role in driving the ecology of venomous snakes (9–11). Although several studies have estimated snakebite risk’ spatial and temporal heterogeneity based on statistical extrapolations and climatic variables, these extrapolations are not based on the processes underlying human-snake encounters dynamics (7, 12–15). Thus, their usage as disease’ monitor tools to improve snakebite management is limited due to the complexity of their extrapolation to countries with no epidemiological data (1, 16, 17).

Epidemiological modeling has emerged as a tool to understand the processes underlying diseases’ spread, and it has been used widely for several NTDs (18–23). Although snakebite envenoming is not an infectious disease, it is caused by the interaction between humans and venomous snakes, and this interaction can be modeled similarly to an infectious disease. A previous study has demonstrated that the main assumption of several epidemiological models (the law of mass action), could explain snakebite geographical variation in Costa Rica (24). By combining models with spatial and temporal surveillance data, it is possible to have reliable estimations for multiple epidemiological parameters and to perform extrapolations of the model to places without data, making these models a useful tool to understand, manage and control several NTDs (25, 26). Nevertheless, no epidemiological models have been used to mechanistically understand the drivers behind the temporal patterns of snakebite, nor to make predictions and estimations of snakebite incidence.

To apply epidemiological modeling to snakebite it is important to understand the biology of the most important venomous snakes’ species, which often is poor or incomplete (10, 27). In neotropics, one of the most important venomous snake groups is the genus *Bothrops* (Wagler, 1824), which is distributed throughout the south of Mexico to the North of Argentina and causes the majority of the envenomings in this area (28–30). This genus has a viviparous strategy of birthing, so each female snake can give birth to several newborns, and, for several species from this genus, the birthing season occurs during the rainy season (9, 11, 31–33). This seasonal dynamic usually increases the venomous snakes’ populations during the rainy season, thus increasing the likelihood of human-snake encounters during this period. On the other hand, rainy seasons can cause flooding, habitat perturbation, and an increase in prey abundance, making venomous snakes more active, resulting in an increase in the likelihood of the encounters too (34– 36). Previous studies have found that snakebite temporal dynamics tend to be related to rainfall seasonal patterns in tropical countries such as Costa Rica and Sri Lanka, but these studies didn’t account for climatic heterogeneity in both countries (13, 14). Therefore, seasonal variation in rainfall has been proposed as an important driver of snakebite incidence temporal pattern, but the possible heterogeneity of this relationship in different climatic scenarios is not well understood.

Colombia becomes a perfect setting to study this association given its location with diverse climatic conditions, where different regions have different rainfall seasonal patterns (37). In the country, snakebite is a severe public health problem, where recent reports account for around 4500 envenoming cases with at least 40 deaths each year, and two species (*Bothrops asper* and *Bothrops atrox*) cause most envenomings (38, 39). In this study, we have developed and calibrated an epidemiological model that can disentangle the role of rainfall on snakebite temporal patterns. We used epidemiological stochastic models in combination with iterated filtering statistical inference methods to disentangle the role of rainfall on snakebite temporal patterns and to understand the geographic distribution of this association.

## RESULTS

In Colombia, between 201 and 422 envenomings are reported every month, with an average of 3659 cases per year (Min: 3135 cases in 2010, Max: 4089 cases in 2015). We found that the number of reported cases has increased with time (Pearson correlation coefficient: 0.58, p-value < 0.05), a result that might be caused by the improving reporting system in the country rather than an increase in the true incidence. To remove this trend from our analyses we used a detrending algorithm based on locally estimated scatterplot smoothing (‘stats’ package, r environment (40, 41)) to only account for seasonality and inter-annual variation in the data (View Figure 1.a for reported and detrended incidence and precipitation data). At a national scale, a cross-wavelet and a cross-correlation function between the detrended national incidence and precipitation show a seasonal correlation between both variables. The estimated seasonality is around 6 and 12 months, and both time series are correlated around a lag equal to zero (View Figure 1.b and Figure 1.c).

**Figure 1.**
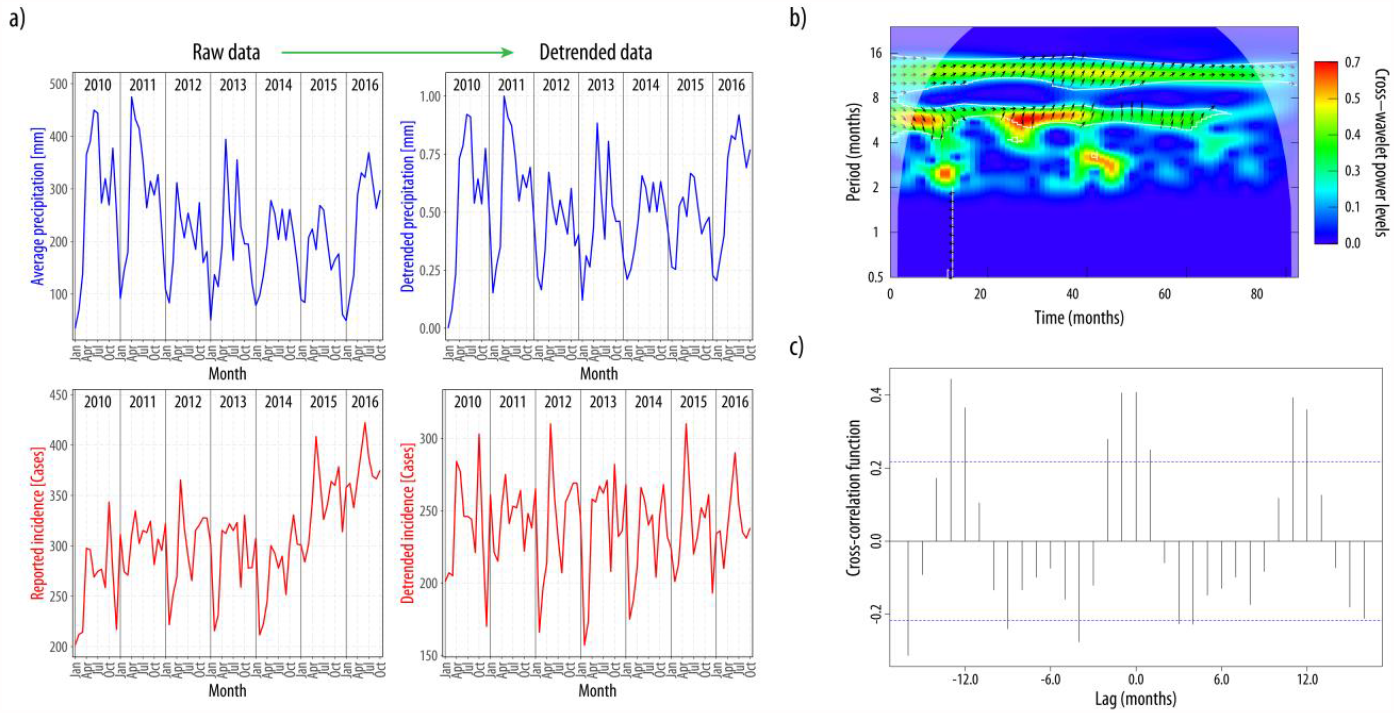
a) Raw and detrended rainfall and snakebite incidence data. To understand the association between rainfall and incidence, we only looked as seasonal and inter-annual variations of both variables, so we worked with detrended variables. **b) Cross-wavelet function between detrended precipitation and incidence for the whole country**. There are 2 significant areas of correlation between both variables that correspond to periods of seasonality of approximately 6 and 12 months. **c) Cross-correlation function between incidence and rainfall for the whole country**. It can be seen that both time series are correlated with no lag, and that they have an annual seasonality (peaks of positive cross-correlation at lags of -12, 0 and 12 years).

To capture the spatial heterogeneity in the rainfall patterns, we clustered the precipitation time series using a k-shape algorithm (42), where an optimum number of 6 clusters was estimated, based on the minimum David-Boulin star index value (0.74) and a silhouette index located in the first quantile of the index distribution (Silhouette index for 6 clusters: 0.46, maximum Silhouette index: 0.5 for 12 clusters) (43). The identified clusters are shown in Figure 2a, defining the regions: *1. South-west, 2. Andean-Pacific, 3. Orinoco-Amazonian piedmont, 4. Central Amazonas, 5. Eastern Orinoco plains, and 6. Caribbean coast*.

**Figure 2.**
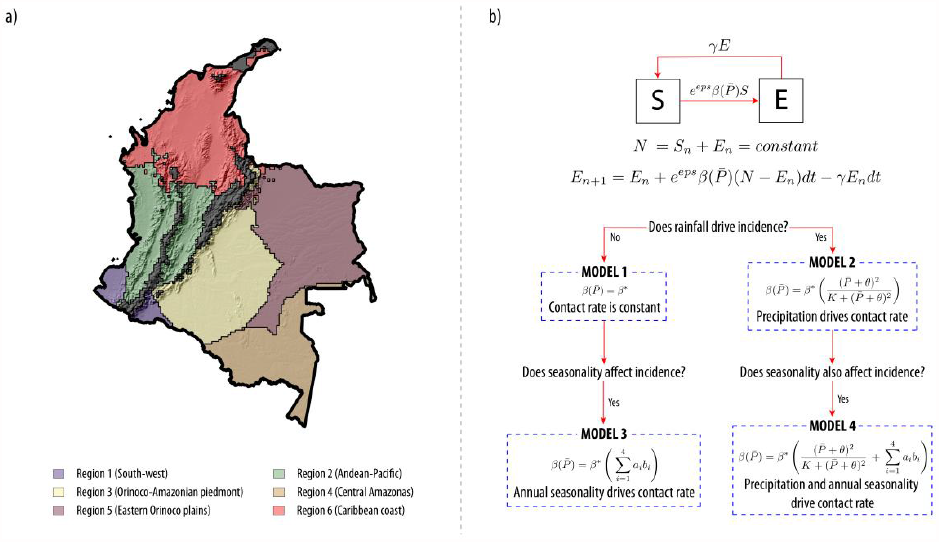
a) Precipitation clusters for Colombia. After the k-shape algorithm, we selected an optimum of 6 clusters because that number of clusters had the best evaluating indexes (DB star=0.74 (minimum value), SIL=0.46 (first quantile, one of the highest values)) for clustering performance. Cluster 1 is the south-west part of Colombia, cluster 2 is the Andean and pacific regions of the country, cluster 3 is the Orinoco-Amazonian piedmont, cluster 4 is the Central-Amazonian region, cluster 5 is the eastern Orinoco plains area, and cluster 6 is the Caribbean coast and Low-Magdalena region. **b) Discrete epidemiological models proposed to estimate snakebite incidence dynamics**. We assume that a susceptible person (*S*) can be bitten by a venomous snake with a probability given by the expression in the arrow between *S* and *E*, where *eps* is a random noise over the encounter frequency with venomous snakes (*β*), and after this bite a person became envenomed (*E*). We assumed no mortality because we want to explain only incidence behavior. Then, an envenomed person can recover with a probability given by the expression in the arrow between *E* and *S* (*γ*), so he becomes susceptible again. For a deeper explanation of the model, see S1) **MODEL 1:** This model doesn’t use precipitation as an input to determine snakebite incidence, so the contact probability *β* doesn’t depend on precipitation (*P*). **MODEL 2:** This model uses precipitation as an input to determine snakebite incidence, so the contact rate *β* depends on precipitation by a type III functional response. **MODEL 3:** This model is used to test seasonality on areas where rainfall does not drive snakebite incidence. Here, we proposed 4 B-splines, each one making one peak each year. First B-spline make peak on October and November, second make peak on January and February, third on April-May, and finally fourth on June and August (View Figure S1.1). In this model, *β* is a linear function of the four B-splines. **MODEL 4:** This model tests if there are seasonality on snakebite incidence that is not explained by rainfall on areas where snakebite incidence depends on precipitation. Here, we summed to the type III functional response the same lineal combination of the same 4 B-splines used on Model 3.

We fitted the proposed stochastic epidemiological model to the national detrended data (Fig 2.b) using an iterative filtering approach that maximizes a likelihood. We tested two different hypotheses by comparing a model that does not account for rainfall as a covariate with a model that accounts for this covariate (Model 1 and Model 2) to answer if rainfall drive snakebite incidence. After fitting models at the national scale, Model 2 was selected as the best model (P-value=0.0062; Table 1), so indeed rainfall drives incidence in the aggregated data. Then, we compared Model 2 and a model that account for an additive seasonality to rainfall (Model 4) to look for additional effects of other seasonal factors, and we found that Model 4 explained better the data than Model 2 (P-value=0.037; Table 1), thus there is another seasonal component in national-aggregated data. The details of the models are shown in S1, and the adjustment for Model 4 at a national scale is shown in Figure S2.

**Table 1.**
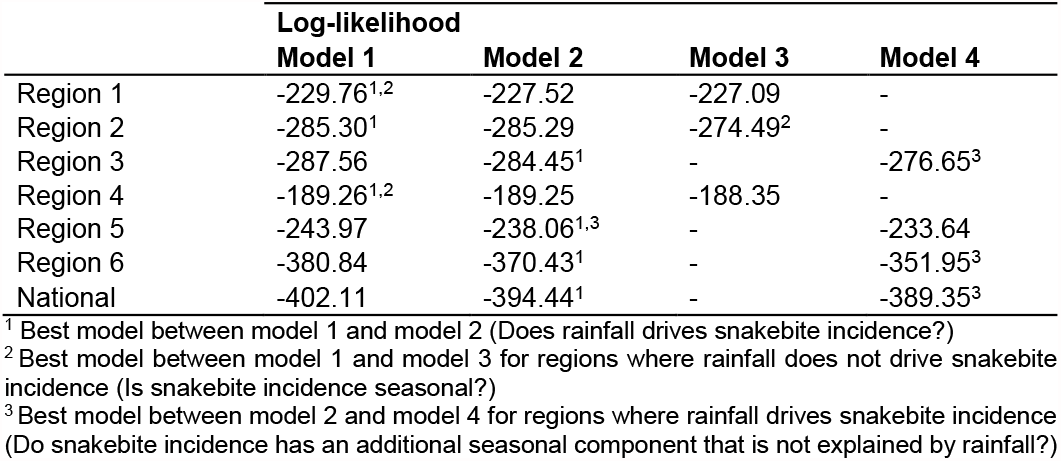
Maximum log-likelihoods for fitted models.

Using the clusters identified previously we fitted our models for every region depicted in Figure 2. a. Interestingly, a diverse pattern of associations between rainfall and incidence throughout the 6 regions of the country was found. We determined that three regions (region 1, 2, and 4) did not have any relationship between snakebite incidence and precipitation. Region 1 and 4 did not exhibit a seasonal component on its incidence (Model 1), while region 2 has a significant seasonal component that is not explained by rainfall (Model 3). On the other hand, region 5 has a strong relationship between incidence and rainfall, but it does not have an association with other seasonal components (Model 2). Finally, the incidence in regions 3 and 6 are associated with precipitation and have other seasonal components (Model 4). The likelihoods of the adjusted models for each region can be seen in Table 1, the likelihood profiles of the parameters for each model are shown in Figure S3, and the confidence intervals for the parameters are shown in Table S4.

We found that the confidence intervals for the parameter K in model 2 (model with rainfall as a covariate) are undetermined in regions where the best model was model 1. In addition, we found that the parameters that account for rainfall in model 4 (K and θ, model with rainfall and seasonality as a covariate) became undetermined as a consequence of the flexibility of the b-splines, which can represent rainfall seasonality (View Table S4). Finally, cases decrease during the dry seasons for regions with the strongest association between rainfall and snakebite incidence (region 5 and 6), where these dry seasons are marked: Both have the minimum monthly rainfall compared with other regions (region 5: 15mm, region 6: 3mm), and this season last at least 3 months (View Figure 3). Thus, rainfall acts as a limiting resource in snakebite dynamics, where regions with marked dry seasons exhibit a relationship between rainfall and incidence mediated by a decrease of cases during drought.

**Figure 3.**
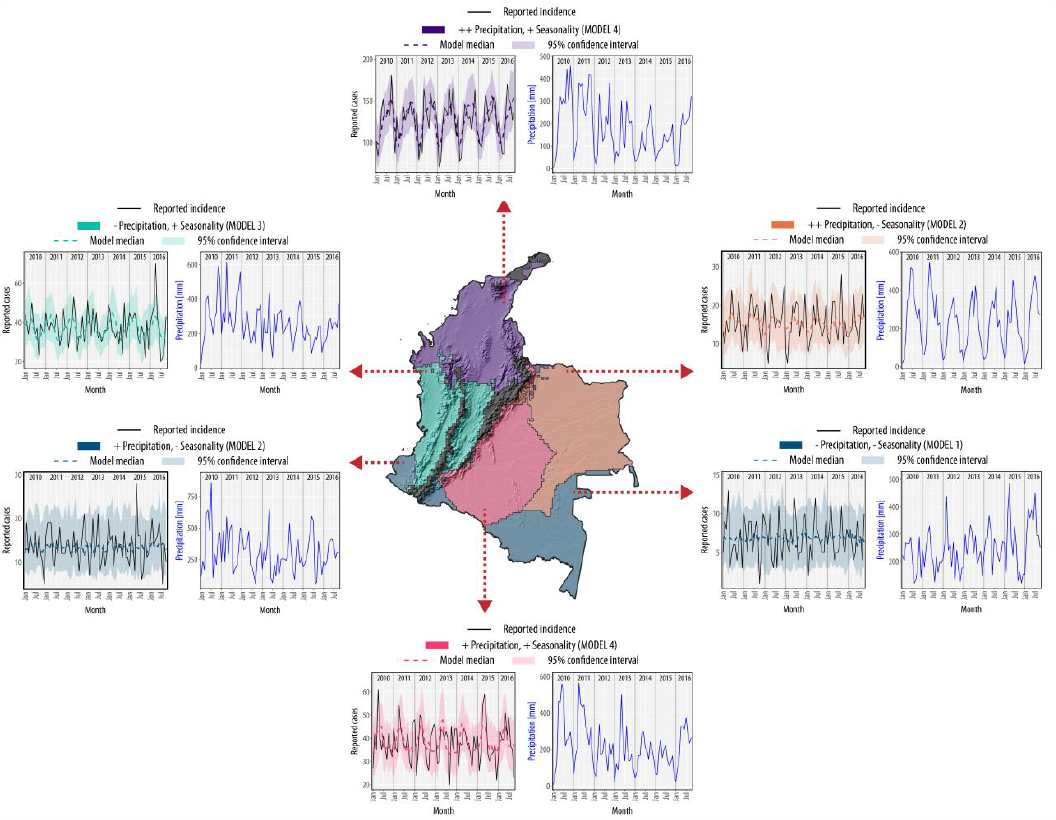
Clusters where snakebite incidence is affected by precipitation, and best-model simulation results. In the map, purple represents regions where we found a strong association between precipitation and incidence with a seasonal component, red is a region with a weak association between rainfall and incidence and a seasonal component, orange is a region with a strong association between both variables and no seasonal component, green is a region where incidence is not driven by rainfall, but has a seasonal component, and bluish gray regions have no association between incidence and both explanatory variables (Rainfall and seasonality). The median of the best model simulation is represented by the dotted line in the time-series plots, while black solid line is the detrended reported incidence, and the ribbon is the 95% confidence interval. We found that precipitation does not drive incidence in 3 regions (regions 1, 2 and 4: *South-*west, *Andean-pacific* and *Central-Amazonas*), that this variable slightly drives envenomings in other region (region 3: *Orinoco-Amazonian piedmont*), and that for the other regions (regions 5 and 6: *Eastern Orinoco plains* and *Caribbean coast and Low-Magdalena*) precipitation drives strongly snakebite incidence. On the other hand, only region 3, and 6, which are regions where snakebite incidence is driven by rainfall, has an additional seasonal component, while region 5 has no additional seasonal component. For regions where snakebite incidence is not associated with precipitation, only in region 2 we found a seasonal component.

## DISCUSSION

Our study quantifies the role of rainfall on the seasonality behind snakebite in different Colombian regions. We found that envenoming seasonality is significantly explained by rainfall at the national level. However, on a finer scale, this relationship is only evident in certain regions with marked seasonal rainfall patterns. The aggregated results suggest that national snakebite incidence has two peaks, the first occurring between April and June, and the second around October (View Figure S2), and the temporal pattern of incidence is driven by rainfall. This association at the national level can be explained by the fact that region 6 (Caribbean coast and Low-Magdalena region) contributes to 54 % of the national cases, and in this region, incidence exhibited a significant association with rainfall (View Figure 3). As a consequence, inferring snakebite dynamics based only on national results should be done carefully: In region 2 (Andean and pacific regions), the peak of envenomings occurs during the beginning of the year, in contrast with the national trend (View Figure 3). Contrary to current studies about snakebite seasonality that have used national-aggregated incidence data (13, 14), our study compares snakebite risk between different regions by analyzing hypotheses represented by epidemiological models: National-aggregated analyses can be largely neglecting the geographical heterogeneity in the association between snakebite and temporal drivers.

### Spatial distribution of snakebite risk in Colombia

By looking at the model without rainfall nor seasonality as covariates (Model 1), the parameter *β* represents a “constant” snakebite risk for each area or an encounter frequency with venomous snakes as described in (24). The likelihood profile over this encounter frequency (View Figure S3.1) and its confidence intervals (View Figure S5) can be used to compare risk between regions: The regions with the highest risk (region 1, 3, 4, and 5) have the presence of *Bothrops atrox* (30, 44), while regions with only *Bothrops asper* (regions 2 and 6) are the regions with the lowest risk. This effect can be caused by ecological differences between both species, which makes *Bothrops atrox* more dangerous or abundant than *Bothrops asper*, or by economical or sociological differences that can increase the exposure to snakebite of inhabitants of these areas. Sadly, biological information about venomous species in the country is scarce, so it is difficult to determine which is the cause of these high-risk areas (10, 27, 30, 44, 45).

### Temporal patterns of snakebite incidence in Colombia

We found that regions with the highest coefficient of variation, which accounts for seasonality, for their precipitation (regions 3, 5 and 6, View Table S6) have their reported incidence driven by rainfall, where incidence decreases during dry seasons (In these regions, model 2 explained better snakebite incidence than model 1). The seasonal rainfall pattern in these regions is also larger than in the other regions (region 1, 2 and, 4, View Figure S7). Thus, a strong seasonality on the rainfall, characterized by marked dry and rainy seasons, determines the association between snakebite incidence and rainfall in Colombia. We propose that the association between snakebite incidence and rainfall in Colombia is mediated via a decrease in the incidence caused by marked dry seasons, which can decrease venomous snakes’ activity, abundance, and finally the encounter frequency between humans and snakes (34, 46).

Another important seasonality measurement is the strength of seasonality, which compares the variance in the noise component of the time series with the variance in the seasonal component (47). Rainfall’s strength of the seasonality was considerably lower in regions where we did not find an association between precipitation and incidence, thus the rainfall signal in these places is noisy and does not have a clear seasonality (View Table S6). It is important to clarify that region 1 has a part located in the Pacific versant, and another is located in the Amazonian versant, which are two distinct ecological regions with different diversity of venomous snakes, but a similar rainfall pattern (30, 48, 49). Thus, these differences in venomous snakes’ diversity and ecology can explain the high noise over incidence time-series in this place. We recommend investigating more deeply the association between rainfall and incidence in this region, where data from Ecuador, which is the neighboring country to this area and has the same species composition in the Pacific and Amazonian versant (44), can help to clarify the dynamics of snakebite incidence in this region.

The mechanisms behind the association between rainfall and snakebite incidence patterns are still unclear, but we want to propose 3 hypotheses to explain this binding: *i)* Several neotropical snakes, including some species of the genus *Bothrops*, have their reproductive cycle related to precipitation pattern: Gravid females give birth neonates at the beginning of the rainy season, thus increasing the abundance of venomous snakes and the probability of encounter between a venomous snake and a human (9, 10). For example, in Costa Rica, the reproductive cycle of *Bothrops asper* is known, and the seasonality of snakebite incidence is driven by the population dynamics of this species (10, 13). This can explain the seasonality found in region 2, and the association between rainfall and incidence with another seasonality component found in region 6. Nevertheless, *Bothrops asper* populations in Colombia are genetically different than populations in Costa Rica, so population dynamics between both populations may vary (50). In addition, it is known that in Brazilian Amazonas the reproductive cycle of *Bothrops atrox* is not so seasonal, where births occur during most of the year (11). This fact can explain why snakebite incidence is not associated with precipitation nor seasonality in region 4 (central Amazonas), but for regions 3 and 5, where *Bothrops atrox* is also present, the incidence is driven by rainfall. *ii)* Precipitation can affect the ecology of venomous snakes, either by causing floods which decrease the area that snakes share with humans, or by increasing ecosystem productivity: More preys (Amphibians, lizards, and rodents) will be available so snakes could be more active thus increasing the probability of a human - venomous snake encounter (10). *iii)* During rainy seasons, agricultural and cattle productivity increases, causing an increase in the number of farmer workers at risk of encounter a venomous snake (51). These three hypotheses are not mutually exclusive, but given that ecological information about venomous snakes is inexistent in Colombia, fieldwork must be done to determine how these three rainfall-related events affect snakebite incidence seasonality and determine risky seasons.

### Final remarks

Our models are capable to estimate the parameters that describe the dynamics of snakebite incidence in Colombia. We believe this modeling framework can be used easily in other countries to monitor snakebite incidence and finally to improve disease’ management under changing environments. It is important to account for regions with different precipitation patterns to determine specifically which snakebite dynamics are happening in the country where this model will be applied. These results demonstrate how countries affected by snakebite can determine in which areas and in which time they need more antivenom, and how to distribute this scarce resource more accurately. This strategic distribution can help to decrease disease’ burden because most places affected by snakebite have a deficit in antivenom coverage (4, 5, 7, 17). In addition, determining the ecological mechanism behind model-estimated snakebite seasonality can help to develop prevention strategies and to capacitate exposed populations to decrease the number of envenomings during risky seasons, thus decreasing the burden of this severe NTD.

We used for the first time an epidemiological model to understand the temporal patterns of snakebite incidence, by using as explanatory variables precipitation and a general seasonal component determined by the B-splines. Given that epidemiological models are robust, and based on the interactions behind the causes of the disease, several modifications and hypotheses can be tested: We encourage researchers to use these epidemiological models to explain and understand snakebite epidemiology. These models have been used in several neglected tropical diseases caused by zoonosis, where the basic biology of the animals involved in disease transmission is known (52– 55). Thanks to this synergy between mathematics and biology, more specific prevention and control programs have been done to decrease the disease burden over the affected population (26, 56–58). Thus, as has been proposed before, the role that the biology of venomous snakes plays behind disease’ epidemiology is important, but it is still neglected: We also want to encourage the study of the natural history and ecology of venomous snakes to fill this enormous vacuum of information that limits the understanding of snakebite epidemiology.

Finally, our work quantifies the association between rainfall, seasonality, and snakebite incidence in Colombia, which is spatially heterogeneous. It is important to determine which ecological or social mechanisms are driving these associations to decrease incidence during risky seasons. With temporal socio-economic data and by using the algorithm developed in this study, novel models can be fitted to quantify the effect of this covariate. Also, these models can incorporate venomous snakes’ population dynamics if the population parameters are measured in the field. Both strategies incorporate the needing of collaborating with social and biological sciences to confront snakebites. Thus, an interdisciplinary approach, with a centralized scope on disease management, must be done to decrease the high burden of this NTD in countries with low healthcare coverage, and finally to help to contribute with the WHO target to reduce snakebite fatality by 30% by the end of 2050 (59).

## METHODOLOGY

We followed three steps to develop and fit an epidemiological model that understands the role of climate, mainly rainfall, on snakebite dynamics. *i)* We proposed four epidemiological models to test the association between rainfall and incidence seasonality, and we fitted them to national data. *ii)* To disaggregate data, we first divided the study region into areas with similar precipitation patterns, so we could aggregate municipality reported the incidence to reduce its noise *iii)* Finally, we fitted our process-based models to an extensive snakebite surveillance dataset by using an iterated filtering approach that maximizes a likelihood (60), and then we explored the role of rainfall in the snakebite dynamics by performing hypothesis testing between different models calibration.

### Epidemiological models

We formulated four discrete epidemiological models to test if precipitation drives snakebite dynamics for each precipitation region. All models assume that incidence is proportional to the number of susceptible populations (*S*) and that the parameter representing this proportionality is *β*: The encounter frequency with venomous snakes. We also implemented a normal random noise to this frequency, which is the parameter *eps*. Additionally, we assumed that each time step a proportion of *γ* of the envenomed population (*E*) recovers from the envenomation (For a detailed explanation, see S1). The first model does not use rainfall as a covariate, but the second model does. In the second model, we established the relation between *β* and rainfall as a type III functional response (View Figure 1.a). Comparing the fit of both models with data will let us test the hypothesis that incidence is affected by rainfall.

To test if there is any seasonality in the incidence in areas where rainfall does not drive incidence dynamics, we proposed the third model, which accounts for seasonality using 4 B-splines, where *β* will be a linear function of four B-splines. These splines are functions that make seasonal peaks on different months of the year: First B-spline makes a peak in October and November, second makes a peak in January and February, third on April-May, and finally fourth on June and August (View Figure S1.1). Finally, model 4 account for precipitation and seasonality, where *β* will be the sum of the ones in Model 1 and Model 2. In addition, this fourth model was only used on clusters where rainfall drives incidence dynamics. The four models used in this study can be seen in Figure 1.a. (View S1 for details of the models). We then fitted each model to the reported incidence data for the whole country and then for every precipitation cluster to determine which model fit better the data and to determine if precipitation and seasonality play a role in driving snakebite incidence dynamics in each area.

#### Fitting epidemiological models to snakebite incidence data

We assumed the process of snakebite envenoming described by epidemiological models as a partially-observed Markov process (POMP), where data is partially observed and we can declare an observation model to estimate the error of data-reporting (60). This observation model was declared as a Poisson process, and the mean of this Poisson model is the monthly flow of people between S and E. This flow is the reported incidence, and it is modeled as the expression shown in the arrow between S and E in Figure 1.a.

Before estimating the parameters with the maximum likelihood, we detrended incidence and precipitation time series by using the algorithm described in (41), and we normalized detrended precipitation between 0 and 1. Then, we estimated the parameters that maximize the likelihood between our model and the data by iterated particle filtering, where we defined a parameter space between the biological limits of the values of the parameters of the models. Then, with a random walk for each point in the parameter space, the algorithm starts to compute the likelihood for each combination of parameters, thus estimating the surface of the likelihood vs the parameters space. Finally, the algorithm converges when the likelihood reaches a “global” maximum (60). The details and settings that we used to perform this estimation can be seen in S8. Finally, we fixed the parameter of recovery fraction *γ* to a value equal to 0.9 for all regions.

After computing the parameters combination that maximizes likelihood nationally, and then per each region (View next section) per each model, we determined if this likelihood were significantly different between model 1 and model 2, so if model 2 outperforms model 1 fitting the data then precipitation explains the dynamics of snakebite incidence in that precipitation area. Then, for regions where model 2 outperformed model 1, we used model 4 to test if there is any additional seasonality in the data. Finally, for regions where model 1 outperformed model 2 (no correlation between rainfall and snakebite incidence), we used model 3 to test if there is any seasonality in the data. To compare likelihoods, we used Wilk’s theorem, which defines a threshold of significance between the difference of likelihoods based on a chi-square distribution. The degrees of freedom of this distribution will be equal to the difference of the number of parameters between compared models, and we used a significance level of 0.05 (60, 61). In our selection, model 1, model 2, and model 4 are nested hypotheses, and model 1 and model 3 too. The fitting and selection of the models, was done using the package pomp in r environment (60).

### Incidence and precipitation data clustering

#### Incidence data

We got public available data of snakebite incidence data between January of 2010 to the end of October of 2016 from the Sistema Nacional de Vigilancia Nacional (SIVIGILA) of Colombia. This dataset is reported in epidemiological weeks, and it details the number of cases that sought medical attention for each municipality (the smallest political unit in the country) per week. We only worked with municipalities that reported snakebite for all of the study years. First, we looked at the epidemiological calendar of Colombia to convert the timescale of the reported cases from epidemiological weeks to months. We aggregated the weekly reported cases for each month. Then, for weeks that overlapped between two months, we distributed the cases by weighting them based on the number of days of the week that belong to each month. After this process, we got a dataset of monthly reported new envenomation cases from 2010 to the end of October 2016 per each municipality of the country.

#### Precipitation data

We got monthly precipitation maps in raster format for Colombia between the year 2010 to the end of October 2016 with a resolution of ∼ 21.23 km^2^ from TerraClimate dataset (http://www.climatologylab.org/terraclimate.html) (62). We removed precipitation data of areas where the two most important venomous snakes’ species in the country are absent by using an altitude threshold of 1900 m.a.s.l. for *Bothrops asper* and of 1500 m.a.s.l. for *Bothrops atrox*, which are the species that cause most envenomings in the country (38, 47).

#### Clustering of areas with similar precipitation pattern

We reduced the resolution of the precipitation dataset by a factor of 12 to perform the clustering algorithm, and we based the clustering on the time series extracted per pixel. We used a k-shape clustering algorithm, using a distance matrix constructed with a shape-based distance (42). We evaluated clustering performance for a predetermined number of clusters between 2 and 10 by using the silhouette (SIL) and Davies-Bouldin star (DB star) indexes, and then we selected the optimal number of clusters by searching the minimum DB star and maximum SIL (43). After this selection, we obtained an optimal number of rainfall regions for Colombia. This algorithm was made by using the package dwtclust in r environment (63).

#### Aggregating incidence and precipitation data for each region

For each cluster, we aggregated rainfall data by computing the average rain for each one of the clustered regions. To aggregate incidence for each cluster, we first computed the reported incidence per area for each municipality for each month. Then, we rasterized this incidence per area to generate maps of monthly incidence per area with the same resolution as the clusters map. Finally, we aggregated this incidence by computing the sum of the incidence per area for each cluster. The outputs of this process are two datasets: *i)* Monthly precipitation for each cluster from the same period, and *ii)* Monthly reported incidence for each precipitation cluster from 2010 to October of 2016.

## CONTACT

Carlos Bravo-Vega

Juan Manuel Cordovez Alvarez

Mauricio Santos-Vega

## COMPETING INTERESTS

The authors declare no conflict of interest.

## DATA SHARING

Data and commented codes to replicate the results of this study are going to be submitted to a data repository suggested by the editorial staff.

## FUNDING

This study was partially funded by Carlos Bravo-Vega: Colciencias, Colombia (https://www.colciencias.gov.co/), Application 727 for doctoral students. The funder had no role in study design, data collection and analysis, decision to publish, or preparation of the manuscript.

## SUPPLEMENTARY MATERIALS

### S1. Epidemiological models and B-splines

We based our models in the law of mass action, where the incidence will be proportional to the contacts between humans and venomous snakes and these contacts will be proportional to the multiplication of the abundance of both populations. This model can be seen in equation 1.

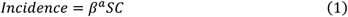

Here, *β*^*a*^ is the absolute contact rate between both populations, *S* is the susceptible population, and *C* will be the venomous snakes population. Given that we don’t know how rainfall affects snakebite incidence, if it is via venomous snakes population dynamics (*C*) or via contact rate (*β*^*^), we merged both in a new parameter *β*, which is what we denote an encounter frequency with venomous snakes. This parameter will depend on a detrended and normalized rainfall 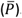. In addition, we added a gaussian noise (*eps*) to this parameter, so our final model for incidence is shown in equation 2.

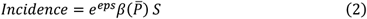

To build the general epidemiological model, we assumed a susceptible population (*S*) and an “envenomed” population (*E*), where the flow from *S* to *E* will be the incidence. We assumed a constant recovery rate for *E* (*γ*), and we assumed no mortality by snakebite. Thus, our general epidemiological model is shown in equation 3.

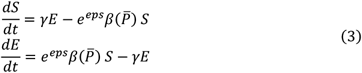

In this model, total population (*N*) will be equal to *S* + E, and it is constant. Thus, we can replace *S* by *N* − E, and we can reduce our system to only one equation that is shown in equation 4.

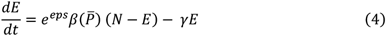

Finally, we can discretize this equation to obtain the final general epidemiological model, which is shown in equation 5.

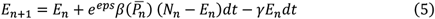

For this general model, we generate 4 models corresponding to different hypothesis over the relation between *β* and *P*. First, for model 1 (Equation 6), the contact rate will be constant. For model 2 (Equation 7), the contact rate will depend on rainfall with a type III functional. Then, model 3 (Equation 8) will assume that *β* depends on only seasonality,

which is a function that depends on 4 B-splines (*b*_*i*_) (View end of S1). Finally, model 4 (Equation 9) will assume that *β* depends on seasonality and rainfall, so it will be a sum of model 3 and model 4.

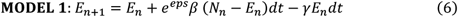

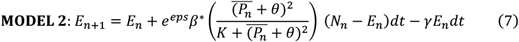

In this model, *β*^*^ is proportionality constant which will set the average value of the contact rate, *θ* is an offset for the detrended and normalized rainfall, and *K* is the constant for the slope of the functional response.

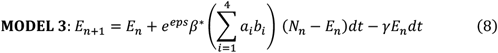

In this model, *a*_*i*_ is a lineal parameter which represent the strength of each seasonal component (*b*_*i*_) in the contact rate.

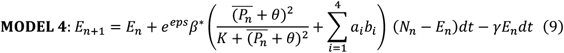

The seasonal components, that depends on the B-splines (*b*_*i*_), can be seen in the figure S1.1.

**Figure S1.1.**
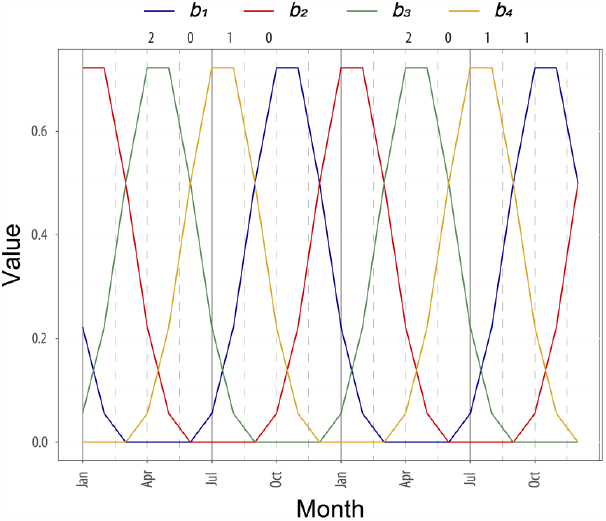
Seasonal components (B-splines) used in the proposed epidemiological models.

**Figure S2.**
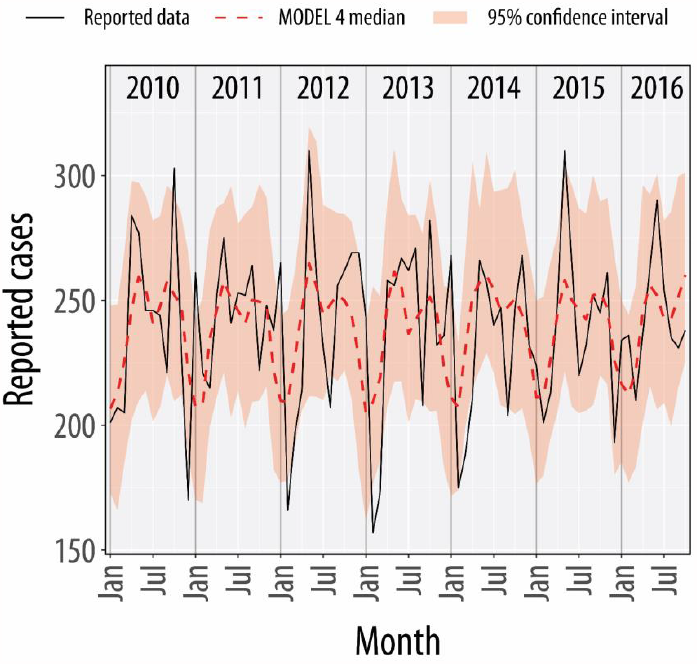
Results for the best model explaining national-aggregated incidence temporal dynamics.

**Figure S3.**
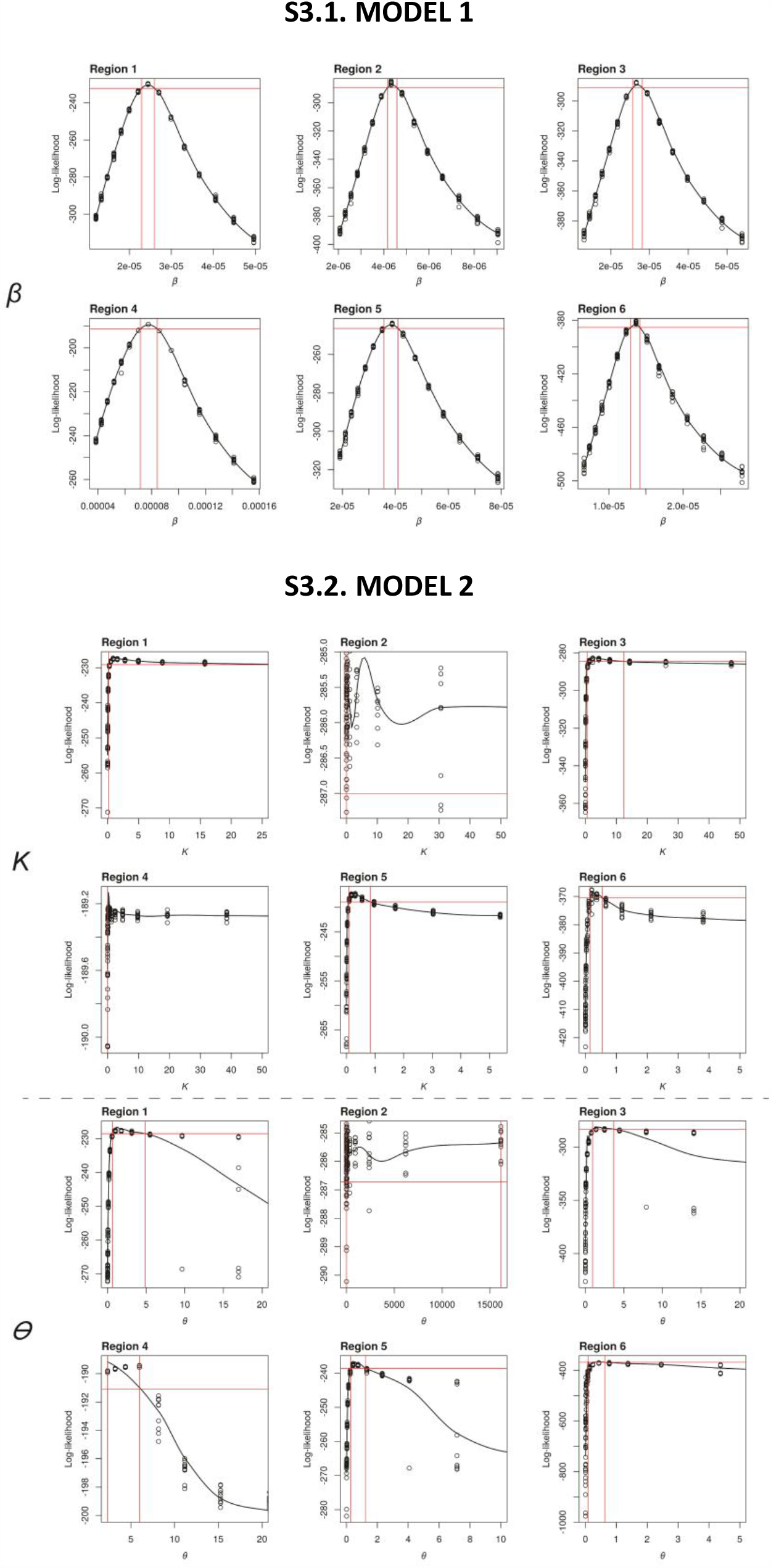
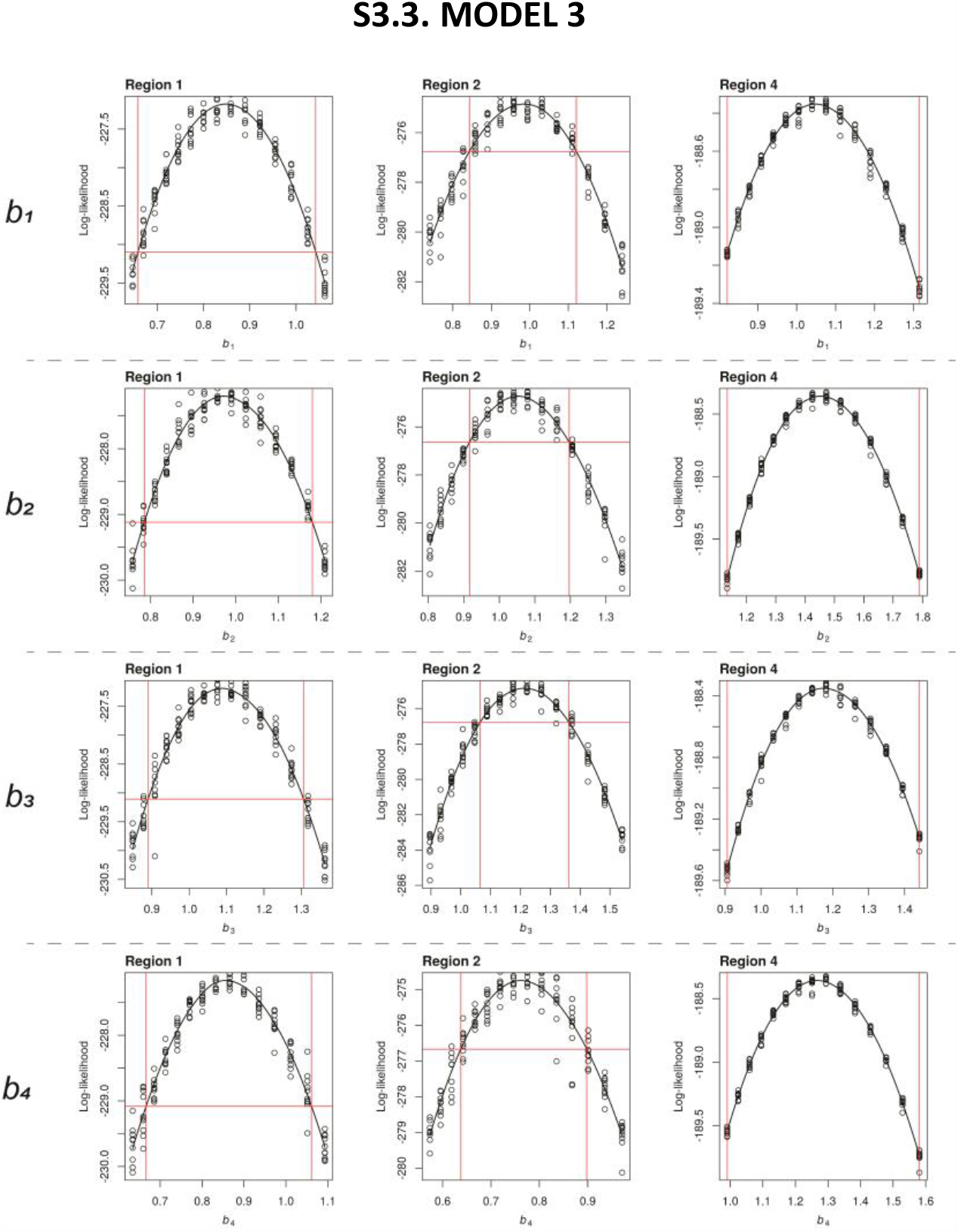
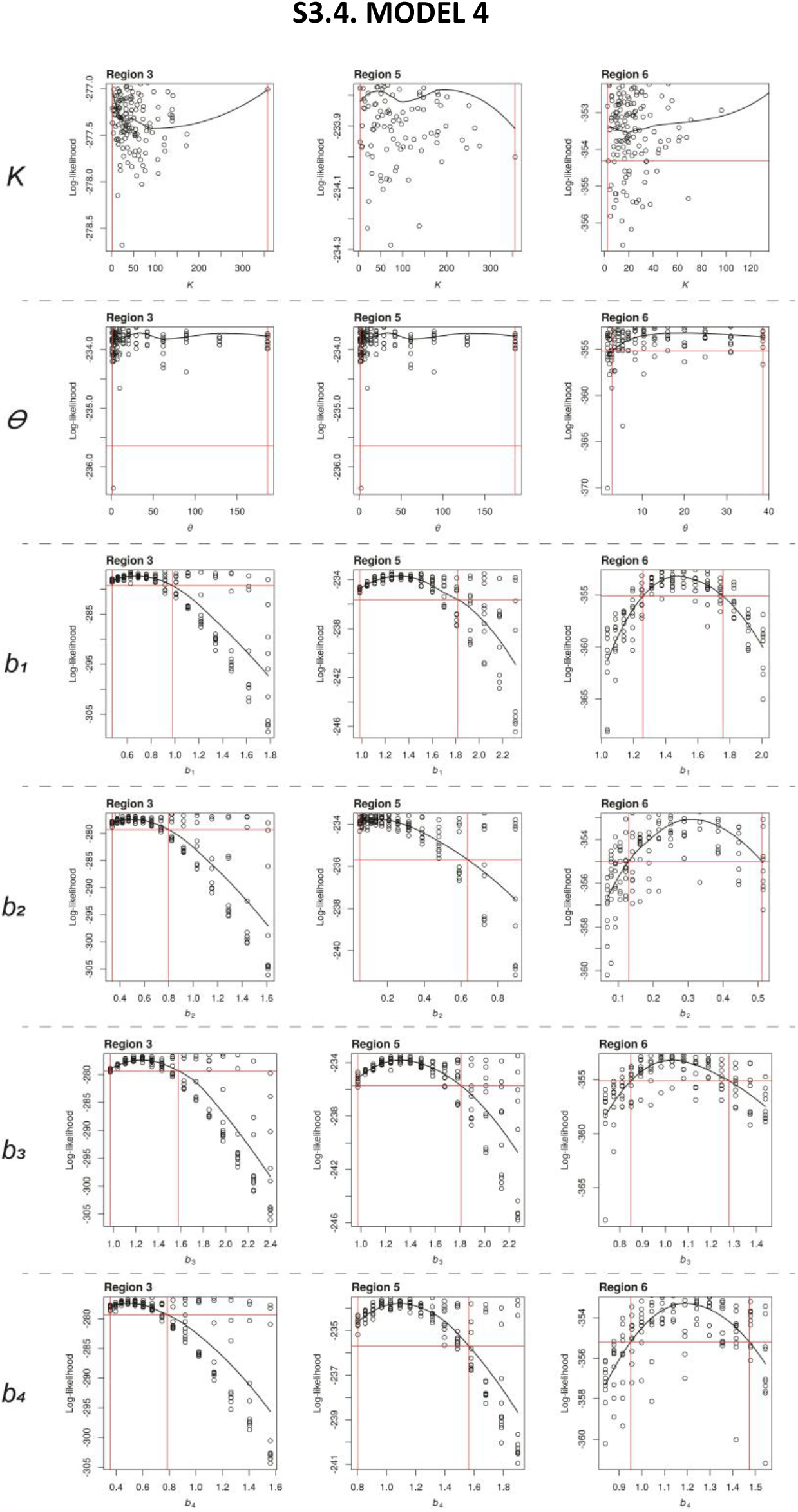
Likelihoods confidence intervals of parameter adjustment for all models in all regions.

**Table S4.**
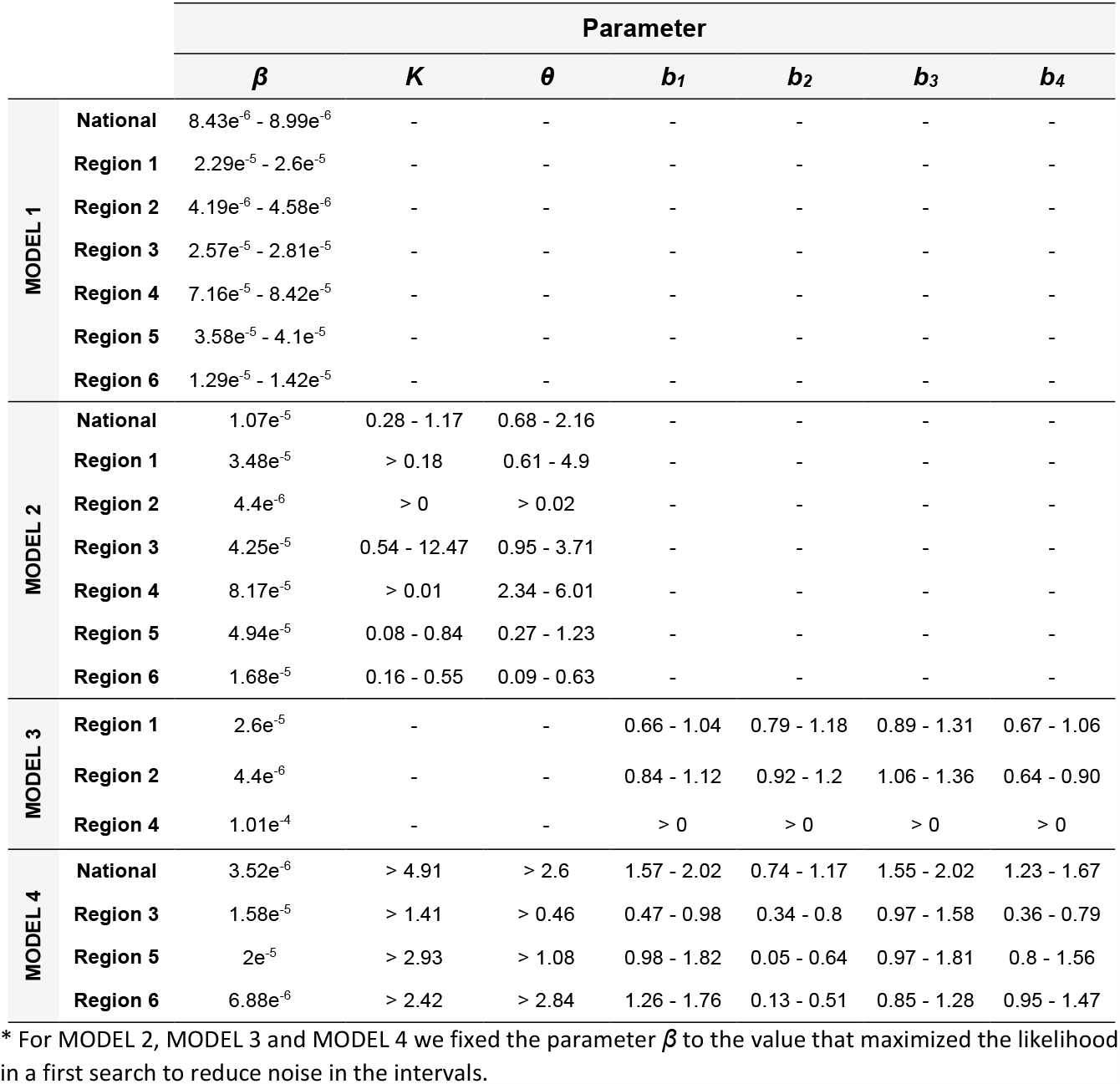
Confidence intervals after parameter adjustment for all models in all regions.

**Figure S5.**
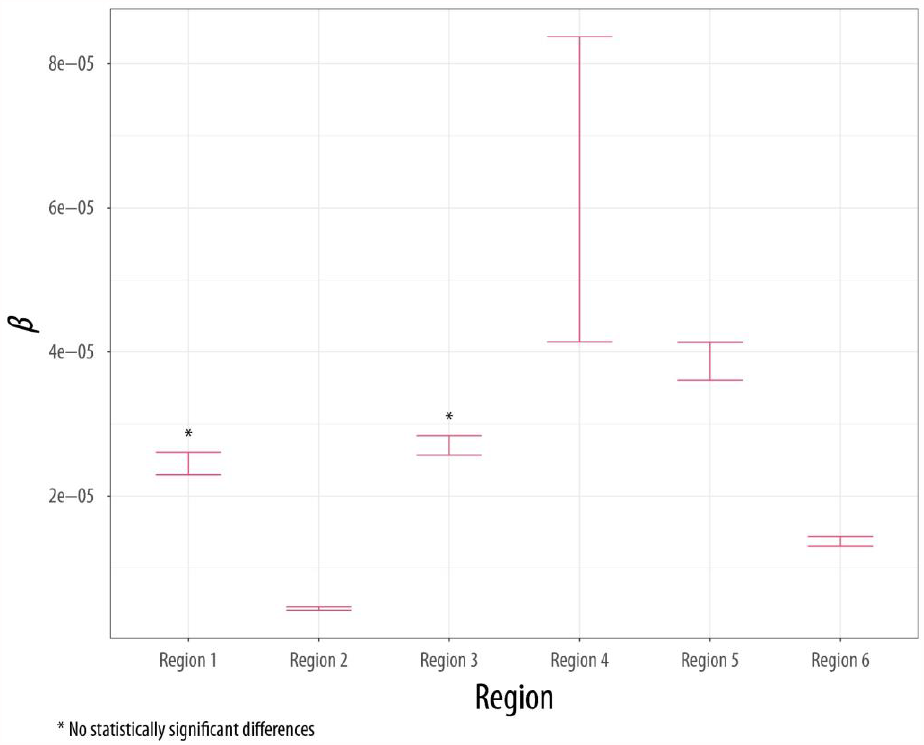
Comparison of the effective contact rate with venomous snakes after fitting MODEL 1 (No rainfall nor seasonality effect) to data.

**Table S6.**
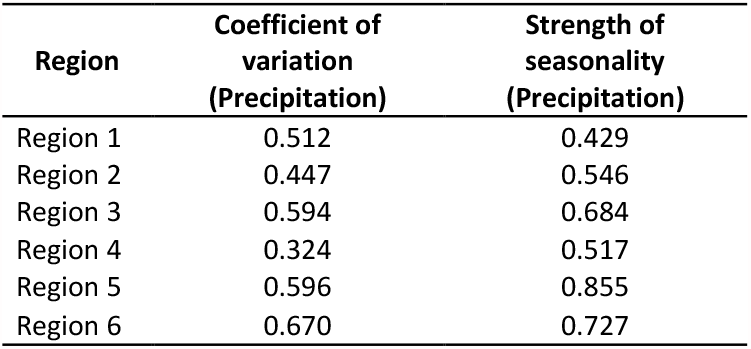
Coefficient of variation and strength of seasonality for precipitation in each region.

**Figure S7.**
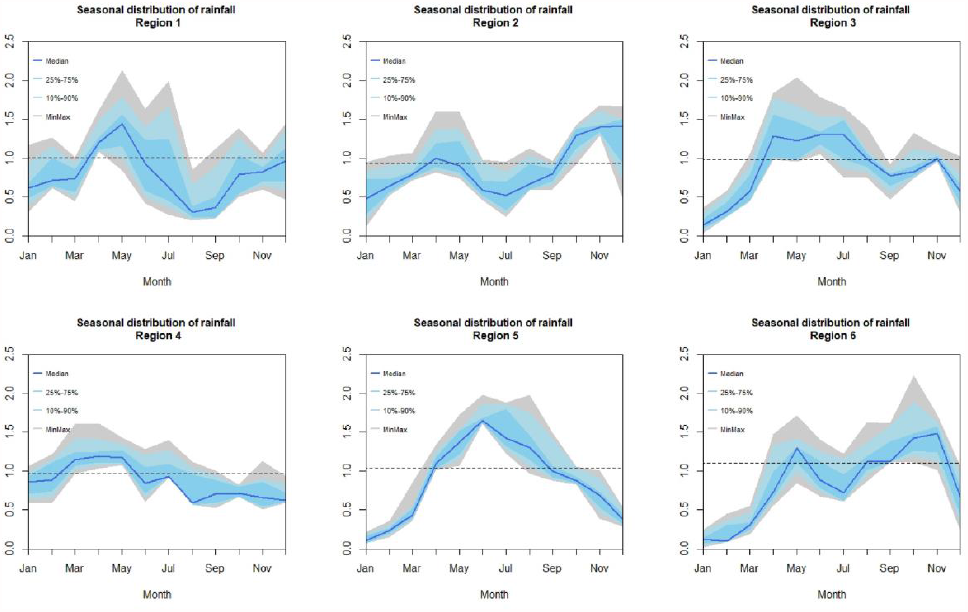
Seasonal distribution of rainfall over different regions of the country.

### S8. Settings for parameter estimation and model selection

We modeled data recording as a Poisson process, with an average value described by the incidence’ model shown in S1. This model is shown in the equation 10.

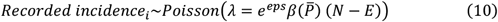

We performed a particle filtering algorithm with 2 steps. The specifications for the first step are:

- Number of particles: 100.
- Number of iterations: 100.
- Magnitude of the random walk perturbations: 0.02.
- Cooling fraction: 0.1.
- Cooling type: Hyperbolic.
- Particles to estimate likelihood for best combination of parameters: 100

Then, over the 10 best fittings for previous step, we performed a second search, defined with these specifications:

- Number of particles: 150.
- Number of iterations: 80.
- Magnitude of the random walk perturbations: 0.05.
- Cooling fraction: 0.1.
- Cooling type: Hyperbolic.
- Particles to estimate likelihood for best combination of parameters: 100

For model 2, 3 and 4 we did both steps to estimate all parameters. Then, we fixed the value for parameter *β*^*^ at the value that maximizes the likelihood to reduce the confidence intervals for the other parameters, and we repeated both steps again.

This work was performed in r environment, using the package ‘pomp’^1^.

^1^King AA, Nguyen D, Ionides EL. Statistical inference for partially observed markov processes via the R package pomp. J Stat Softw [Internet]. 2016 Mar 25 [cited 2020 Mar 30];69:1–43. Available from: http://arxiv.org/abs/1509.00503

## Notes

### Competing Interest Statement

The authors have declared no competing interest.

